# Connectome-based predictive modelling predicts frailty levels in older adults

**DOI:** 10.1101/2025.08.26.672253

**Authors:** Amin Ghaffari, Majd abouzaki, Yasmine Romero, Andrew Sun, Aaron Seitz, Jason Langley, Ilana J. Bennett, Xiaoping Hu

**Affiliations:** Department of Bioengineering, University of California Riverside, CA; Department of cell, molecular, and developmental biology, University of California Riverside, CA; Department of Biochemistry, University of California Riverside, CA; Department of Psychology, Northeastern University, Boston, USA; Department of Psychology, University of California Riverside, CA; Center for Advanced Neuroimaging, University of California Riverside, CA

**Keywords:** Connectome, frailty, task-based functional connectivity, connectome-based predictive modelling, caudate nucleus

## Abstract

Frailty is characterized by a persistent and progressive decline in physiological reserves, leading to increased vulnerability to stressors and a heightened risk of adverse health outcomes, both physically and mentally. Despite frailty’s prevalence in older adults, there is limited research on its neural substrates, especially using task-based brain functional connectivity. In this study, we used connectome-based predictive modelling (CPM) to find a linear relationship between task-based connectomes — taken from tasks that involved similar handgrip manipulations — and a separate measure of frailty: the maximum grip strength in older adults. We observed that the task-based connectomes were able to explain individual differences in grip strength, with the Subcortical and Cerebellum network, particularly the caudate nucleus, functional connectivity being the strongest predictor. These findings demonstrate that task-based functional connectomes can serve as personalized markers that can predict individual behavioral measures, including handgrip strength, and point to involvement of the caudate nucleus in frailty.

**Key points:**

- We used connectome-based predictive modeling on task-based fMRI to predict grip strength in older adults, a key marker of physical frailty.
- The model significantly explained inter-individual differences in contraction strength using functional connectivity patterns.
- Subcortical regions, especially the caudate nucleus, played a major role in prediction, highlighting their relevance in frailty assessment and potential interventions.

## 1. Introduction

Physical frailty, which refers to a decline in physical strength and energy, is prevalent in older adults and has been attributed to impaired cognitive function and adverse health outcomes [1, 2]. The strength of a contraction on a handgrip, known as isometric handgrip strength, has been used as a measure of frailty [3, 4]. While handgrip strength can partially be explained by muscle properties (e.g., cross-sectional area and architecture) [5, 6], it may also be influenced by neural adaptations, such as intermuscular and intramuscular coordination. Moreover, the brain likely plays a crucial role in regulating contraction force production and coordination [7]. These neural contributions to handgrip strength are understudied but may serve as novel biomarkers of physical frailty in older adults, potentially at the individual level.

There is a growing use of imaging-derived data from different modalities to predict clinical phenotypes and disease risk [8-11]. In this context, handgrip strength has been attributed to resting-state functional connectivity within motor and salience networks. For example, within healthy older adults, Seidler and Rachel [12] found that higher functional connectivity of the motor cortex to putamen, insula, and cerebellum was associated with higher handgrip strength. Another study investigated whole-brain functional connectivity (i.e., connectome) and observed that higher handgrip strength was associated with greater functional segregation of the salience ventral attention network in older adults at rest [13]. This highlights the potential of the connectome as a valuable resource for analyzing the neural mechanisms underlying grip strength and frailty in aging.

Because the connectome is unique for each person, akin to a “brain fingerprint” [14], it may also serve as a personalized marker that can be used to predict their individual behavioral measures [15, 16], including handgrip strength. Brain fingerprints serve as the foundation of connectome-based predictive modeling (CPM), a data-driven approach that maps individualized functional connectivity patterns to behavioral phenotypes, enabling the prediction of behavior with personalized precision [17]. Researchers have used CPM with either resting-state or task-based connectomes to explain individual differences in cognitive performance (e.g., intelligence [14], mnemonic discrimination [18], working memory [19]) and clinical symptoms (e.g., anxiety levels [20], internet addiction symptomatology [21], autism symptom severity [22], compulsion severity in obsessive-compulsive disorder [23], depressed and elevated mood severity in bipolar disorder [24],). CPM has also been utilized to predict behavioral measures in older adults (e.g., attentional control [25], trust propensity [26]), demonstrating the viability of this method for phenotypic prediction of this cohort. Extending this approach to frailty, Zúñiga et al. [27] applied CPM to predict self-reported frailty indices in older adults using their resting-state connectomes.

While both resting-state and task-based connectomes could be used in CPM, task-based connectomes may provide a more robust characterization of brain–behavior relationships [28-30]. This is because the brain state is manipulated based on the task being performed [30], shifting the brain connectivity pattern into a more task-relevant state that could be more informative for task-related behavioral measures. Given this closer brain– phenotype coupling, it is important to evaluate whether task-based connectomes derived from tasks sharing similar components maintain individual specificity and predictive power. Such validation is critical for ensuring the generalizability and extendibility of CPM across different task conditions.

To better understand the neurobiological basis of frailty in older adults, a CPM-based predictive model can be developed using connectomes from tasks combining motor and perceptual components. Such tasks shift the brain into a more alert, motor-relevant state, offering greater sensitivity to motor-related conditions like frailty compared to resting-state connectivity. In this study, we focused on healthy older adults and had them perform two perceptual discrimination tasks on two different fMRI sessions, both of which involved a non-dominant handgrip manipulation. We aimed to test the identifiability of the task-based connectomes across the sessions and identify the key functional connections (FCs) that provided this identification power, which we predicted would be derived from the shared handgrip component. We also measured participants’ maximum isometric voluntary contraction (MVIC) of their non-dominant hand as an indicator of frailty and neuromuscular health [31-34], and evaluated the task-based CPM’s ability to predict MVIC in older adults. We identified FCs predictive of MVIC, which may partly explain motor-related impairments in frail older adults. Finding such brain-based biomarkers for grip strength could identify potential target sites for motor rehabilitation programs, enhancing the effectiveness of interventions such as Brain-Computer Interfaces (BCIs) and Functional Electrical Stimulation (FES) aimed at mitigating frailty in older adults [35, 36].

## 2. Materials and Methods

### Participants

Fifty-five older adults recruited from communities near University of California, Riverside (UCR) aged between 65 and 87 years (29 females, M_age_=68.8 yrs, SD_age_=5.8 yrs) participated in this study. Inclusion criteria required participants to be free of MRI contraindications (e.g., non-compliant implants, claustrophobia), have no significant health problems, no history of drug or alcohol abuse, no hearing or uncorrected vision loss, no psychiatric or neurological condition, right-handedness, and not take psychotropic medications. The participants were screened for normal cognition using the telephone version of the Montreal Cognitive Assessment, MoCA. All participants gave informed consent in accordance with the institutional review board at UCR and received financial compensation for their participation.

### Isometric Handgrip Contraction

At the beginning of each testing session, participants were asked to squeeze a digital hand dynamometer (Vernier Software & Technology, Beaverton, OR) at maximum power with their left (non-dominant) hand. They repeated this 3 times, and the values were averaged to calculate their MVIC of that session. The final MVIC for each participant was the average of their MVIC values across the testing sessions. This was the parameter of interest to be predicted using the functional connectome.

Before starting the first fMRI run in each testing session, the MVIC of each participant was measured again inside the scanner using a fiber optic MRI dynamometer from Current Design, Philadelphia, PA. The procedure for this measurement was the same as that of the outside-scanner MVIC. The inside-scanner MVIC was recorded to determine the values required for the participants to perform high grip (HG) and low grip (LG) trials during the fMRI runs. It should be noted that the inside-scanner dynamometer was not linear across the entire range of force, so the outside-scanner one was used to compare MVIC across the participants.

### Image Acquisition

Following the MVIC measurement in each testing session, participants went into a Siemens 3T Prisma (Siemens Medical Solutions, Malvern, PA) scanner at the UCR Center for advanced neuroimaging and MR images were acquired using a 64-channel receive-only head/neck coil (or a 32-channel receive-only head coil if the participants did not fit in the 64-channel head/neck coil). A T_1_-weighted MPRAGE scan (repetition time (TR)/echo time (TE)/inversion time = 2400/2.72/1060 ms, field of view (FoV) = 256 × 240 mm^2^, 208 slices, 0.8 mm^3^ isotropic resolution, GRAPPA = 2) was acquired to be used as the reference for registration of the functional scans to the anatomical space and then the Montreal Neurological Institute (MNI) space. Five runs of a gradient echo-planar imaging (EPI) sequence (TR/TE = 1750/32 ms, FoV = 221 × 190 mm^2^, 72 slices, 1.7mm^3^ isotropic resolution, GRAPPA=2, multiband factor = 3, 7.5 min, 245 TRs) measured the Blood Oxygen Level Dependent (BOLD) signal while participants performed either the auditory or visual task. Additionally, two spin-echo EPI scans were collected (TR/TE = 7700ms /58 ms, FOV = 221 × 190 mm^2^, 72 slices, 1.7mm^3^ isotropic resolution) with anterior-to-posterior and posterior-to-anterior phase encoding directions for susceptibility distortion correction.

### Task Paradigm

On each of the fMRI testing sessions, participants completed either the auditory (ADT) or visual (VDT) detection task (see Figure 1), with a counterbalanced order across the participants. Each task session consisted of five fMRI runs, each with 30 event-related trials. Each trial started with participants squeezing a dynamometer for 3000 ms at either 5% (low-grip, LG; 50% of trials) or 40% (high-grip, HG; 50% of trials) of their in-scanner MVIC. This manipulation was intended to modulate arousal [37, 38], but is of interest in the current analysis because it involved a handgrip demand on each trial. The squeeze was followed by a blank screen (250 ms), fixation dot (500 ms), task-specific initial stimulus (100 ms), fixation dot (500 ms), task-specific second stimulus (100 ms), fixation dot (250 ms), a prompt to respond whether the stimuli were the “same” or “different” (3000 ms), and then a prompt to indicate confidence in the prior response (3000 ms). In the ADT task, the initial stimulus was always a standard tone (1000 Hz). The second stimulus was either a standard tone or an oddball tone that was offset from the standard by +4, +8, +32, or +64 Hz (6 trials each per run). In the VDT task, the initial stimulus was always a standard visual Gabor patch (6 cycles per degree, 4-degree diameter; Michelson contrast of 0.2 between the maximum and minimum luminance values of the Gabor patch). The second stimulus was either the standard Gabor patch or an oddball Gabor that was offset from the standard Gabor’s contrast by 0.04, 0.08, 0.16, or 0.32 Michelson contrast (6 trials each per run). Average accuracy, measured as the percent of correct same/different responses to the second stimulus, was calculated for each contrast level and each task. Confidence ratings were not considered in the current study.

**Figure 1.**
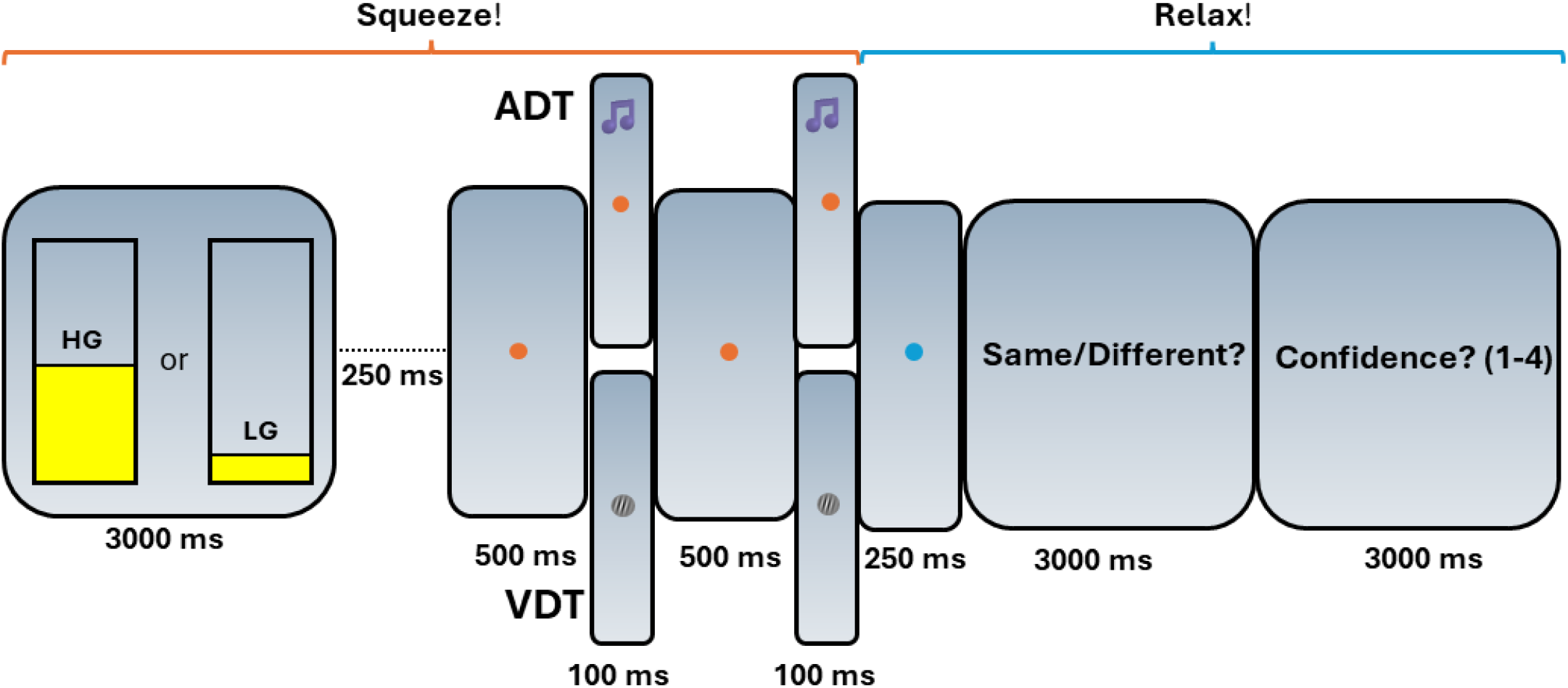
Schematic of the task paradigm. The auditory (ADT) and visual (VDT) tasks share a similar trial structure, differing only in stimuli— two visual Gabors for VDT and two tones for ADT. Participants perform a high (HG) or low grip (LG) squeeze on a dynamometer, maintaining it until the second stimulus ends. They then relax, indicate whether the stimuli were the same, and rate their confidence. Block durations are provided in milliseconds.

### Preprocessing

Raw fMRI and T1-weighted dicom files were converted to niftii files using *dcm2niix [39]*. Then, the T1-weighted structural scans of the participants were skull-stripped using *BET* from FMRIB Software Library (FSL, www.fmrib.ox.ac.uk/fsl). The skull-stripped brain was segmented into white matter (WM), gray matter (GM), and the cerebrospinal fluid (CSF) using FSL’s FAST and the corresponding masks were extracted. The fMRI images were corrected for slice timing, motion, and susceptibility distortions using FSL’s *slicetimer, mcflirt*, and *topup*, respectively. Spatial smoothing was performed using a Gaussian filter with a Full Width at Half Maximum of 2mm.

A transformation was derived from the average functional preprocessed scan of each participant to their T_1_-weighted image using a rigid body transform with a boundary-based registration cost function. Next, a transformation was derived from each participant’s T_1_-weighted image to standard space using a non-linear transformation. The two transformations were concatenated, and each fMRI series was transferred to the MNI space using the resulting transformation. Transformed images were visually examined to ensure accurate registration.

### Functional Connectomes

The Shen et al. 268 region atlas [40], was used to define regions of interest (ROIs) in MNI space (Figure 2a). This parcellation’s regions are grouped into eight functional networks: Medial Frontal (MF), Frontoparietal (FP), Default Mode (DM), Subcortical and Cerebellum (SC), Motor (M), Visual I (VI), Visual II (VII), and Visual Association (VA). VI and VII were combined into a single visual network (VIS) because some ROIs in VII were outside the field of view in this study. Additional ROIs, particularly in the cerebellum, were also excluded, resulting in a parcellation of 215 ROIs across seven functional networks.

**Figure 2.**
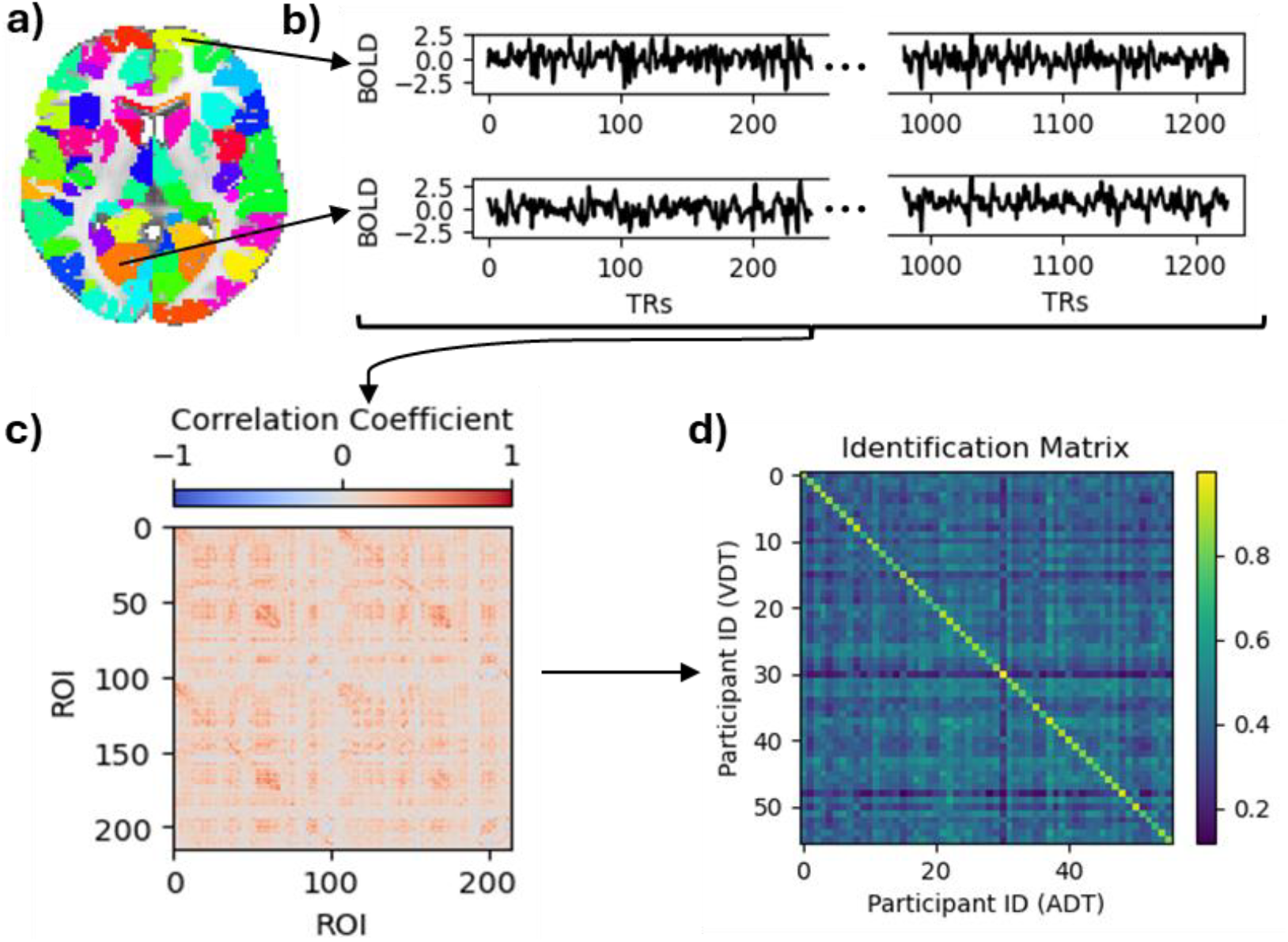
Pipeline of deriving participant-based connectomes and identification. **(a)** An axial view of the parcellation. **(b)** Time courses of the regions indicated with black arrows normalized and concatenated across the runs. **(c)** A sample subject’s connectome of the ADT. **(d)** Identification matrix. The diagonal elements represent the correlation between the same participants’ connectomes across the two task sessions.

For each run, the voxels’ time series within each ROI (node) were averaged to extract the average time series of that region. At this stage, 6 motion parameters (3 translations and 3 rotations) as well as the average signals of the white matter, cerebrospinal fluid, and the global signal were regressed out of each ROI’s time series. Linear trends were also removed from each ROI’s time series and a low-pass filter (0.2Hz) was applied. Then, the time series were z-scored and concatenated across the runs, so each participant had a time series of 245×5 for each of their ROIs (Figure 2b). Pearson’s correlation coefficient was measured between each pair of the ROIs’ time series to produce the participant-specific connectomes (Figure 2c). By only considering the lower triangular part of the functional connectivity maps and applying Fisher’s r to z transformation to the resulting vector, the functional brain fingerprint of each participant was acquired.

### Identification

To observe the performance of the fingerprints in identifying the participants, an identification matrix was calculated by computing Pearson’s correlation coefficient between each pair of the participants’ connectomes across the two sessions (Figure 2d). In this matrix, each row stores a participant’s VDT fingerprint correlation coefficient with all participants’ ADT fingerprints, and if the maximum of the row is the diagonal element, it indicates a correct ADT identification. The same goes for identifying VDT scans within each column using ADT data. We also performed a permutation test with 1000 iterations, in which participants’ identifiers for one task were shuffled. Identification accuracy was then calculated for each iteration, and the resulting distribution (chance distribution) was compared to the accuracy from the real data.

Task-based connectomes are made up of FCs between different ROIs, and each of these FCs (i.e., features) contribute differently to the identification power of the connectomes. We used Differential Power (DP) to measure how discriminative (differentiating) each feature was among participants [14]. To measure DP, we computed the feature product value (FPV) between each pair of the participants across the ADT and VDT session for each feature,

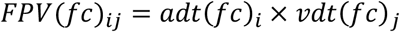

where i and j are labels of the participants, fc represents a FC (feature or edge), and adt/vdt are the functional connectomes of the corresponding participants derived from ADT/VDT tasks. If an FC is discriminative between participants, its FPV when calculated for the same person should have a higher value than for different participants. In other words, a differentiating edge across the functional connectomes has the property that,

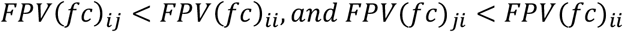

Following the same steps described in previous work [14], we computed the empirical probability (p_e_) of each feature in maintaining this characteristic in the dataset,

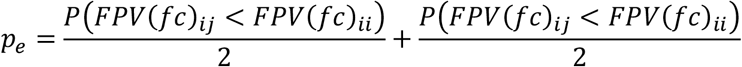

The probabilities in this equation can be calculated by counting the number of participants for whom this characteristic is being held. Finally, DP is calculated by computing the log likelihood of the empirical probability across the dataset.

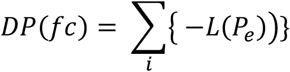

A high DP value is associated with a low value of the p_e_ parameter, meaning that the corresponding FC is more discriminative, and therefore, has contributed more to participants correct identification.

### MVIC prediction

The ability of functional connectomes to predict average (out-of-scanner) MVIC scores was evaluated using 52 participants, as three were excluded due to missing MVIC scores. We used CPM to predict these scores using connectomes of both tasks. In summary, this method consists of one feature selection and one prediction step in a leave-one-out manner [17]. In doing so, first, a subject is set as the test set and the rest are set as the training set. The training set goes into the feature selection step, where the correlation coefficient of each feature of the connectome with MVIC scores across the training set is computed. If the p-value of this correlation coefficient is lower than a certain feature selection threshold (FST), that feature is retained for model building. The retained features are separated into two sets of positive and negative features based on the sign of their correlation with the MVIC scores. Then, for each participant, the value of all negative and positive retained features is averaged separately so each participant has a negative and positive feature strength in the training set. Finally, using the difference between negative strength and positive strength values, a linear model is built to predict the MVIC scores of the training set (each participant has only one entry into the model, that is, their strength difference). This model is then applied to the test set and the test participant’s MVIC score is predicted using their strength difference. By doing this procedure for all participants, a vector of the predicted MVIC scores is generated. The performance of the CPM in predicting the MVIC scores is assessed by measuring the correlation coefficient between the predicted and observed scores.

MVIC scores are known to be affected by age and sex of the participants, where males usually have a higher score compared to females [41, 42]. For this reason, we considered age and sex as covariates and regressed out their effects from the MVIC scores using a linear model. It should be noted that if the data from the test set is used for covariate regression, it might cause inflated performance of the CPM due to data leakage [43]. Therefore, in each fold, we derived a model to regress out the covariates effect from the training set and applied that model on the test participant to remove the covariates’ effect from the test participant. To ensure CPM performance was independent of the chosen FST value, we tested four different thresholds: 0.01, 0.005, 0.001, and 0.0005. To identify the FCs predictive of MVIC and their potential role in frailty, we extracted features consistently retained across all folds for both tasks. We then visualized shared features between the two tasks to ensure the shared features were not task-specific or influenced by outliers.

Permutation testing was conducted for MVIC prediction by randomly assigning MVIC values to brain fingerprints. The CPM with covariate regression was applied to each permuted dataset, independently performing covariate removal, feature selection, and prediction in each iteration. By computing the correlation between predicted and observed MVIC values for 1000 permutations, a chance-level CPM performance was obtained and compared to the CPM performance on real data.

## 3. Results

### MVIC performance

While the partial correlation between chronological age and grip strength—controlling for sex—was not statistically significant in our sample (r = –0.10, p = 0.46), we regressed out age effects from grip strength to maintain consistency with prior studies [44] (Figure 3a). Regarding sex, between-group t-test revealed that males (34,859.49) had significantly higher average MVIC values than females (22347.20), p<0.001.

**Figure 3.**
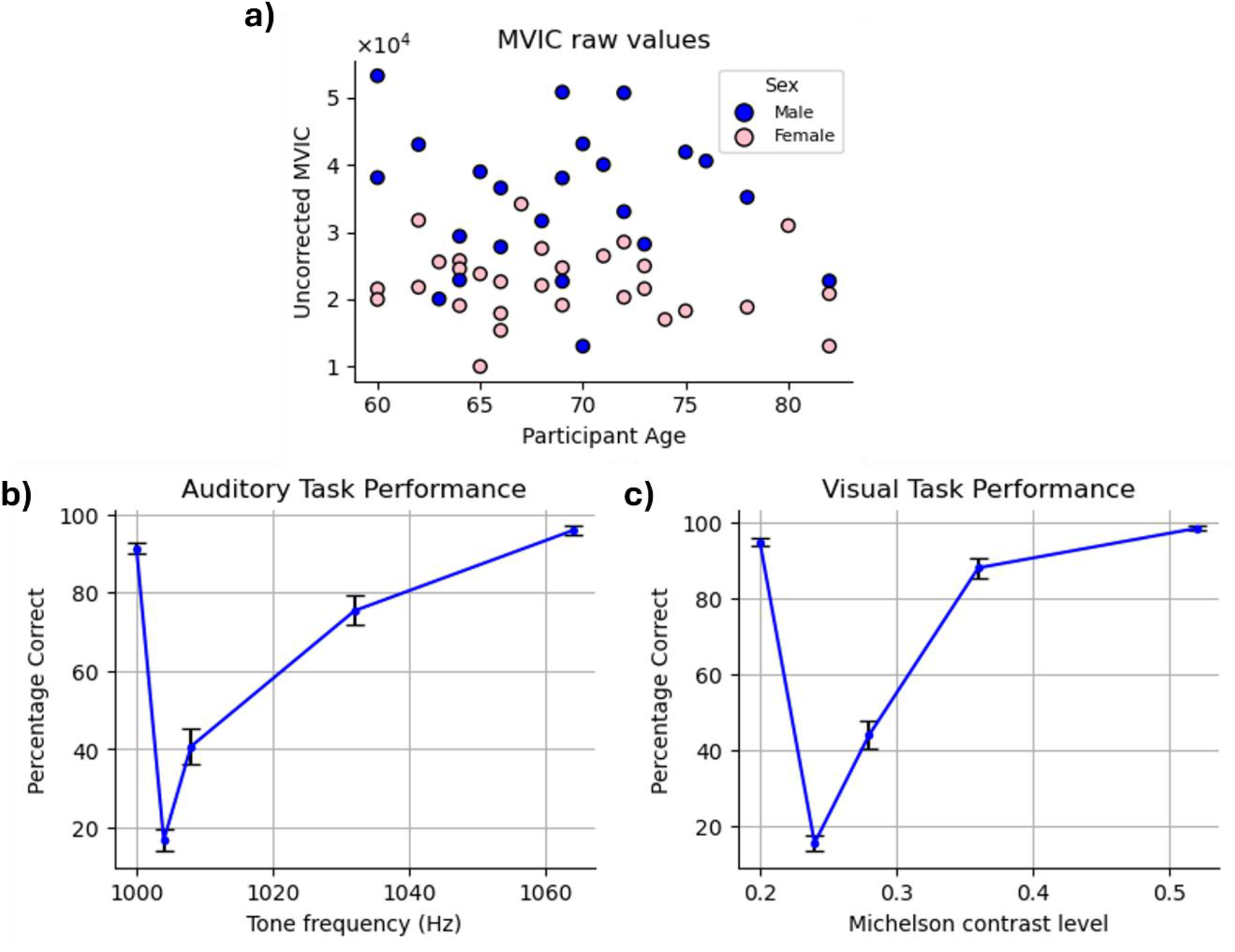
Behavioral Task performance. **(a)** MVIC values. The blue and pink circles in the scatter plot correspond to males and females, respectively. MVIC values are reported in an arbitrary unit reported by the dynamometer. **(b and c)** Average accuracy and confidence ratings across the participants in **(b)** auditory task **(c)** visual task. The red plots show the confidence ratings trend, while the blue ones show the average accuracy. Error bars indicate the standard error of the mean (SEM) across participants.

### Task performance

Examination of accuracy data confirmed that participants exhibited high task performance on both the VDT and ADT, as reflected in their response accuracy. Correct “same” responses to standard stimuli were high (VDT: 92.62%; ADT: 89.29%) and correct “different” responses to oddball stimuli increased monotonically as a function of their dissimilarly to the standard stimulus (i.e., as they became easier to distinguish from the standard stimulus) (Figure 3 b-c).

### Identification

For both VDT and ADT, all participants were correctly identified using the fingerprints derived from the other task (identification accuracy = 100%), showing that the task-derived connectomes are highly reliable as individual-specific markers within this cohort. The permutation test confirmed that this result is not likely due to chance (p < 0.0001).

To recognize the FCs with a significant role in the identifiability of the connectomes and their corresponding ROIs, DP of all the FCs was computed and thresholded at 99.5^th^ percentile to visualize the most discriminative edges and their corresponding nodes. The SC network had the most significant role in providing the differentiating edges for the connectome. Approximately 67% of these discriminative features connected the ROIs of the SC network to each other both within each hemisphere and across the hemispheres (Figure 4 a-b). The FCs between the SC and M network also played an important role in identifying the participants, as about 20% of the suprathreshold DP edges connected ROIs of these two networks (Figure 4a), having both inter- and intra-hemispheric edges (Figure 4 b-c).

**Figure 4.**
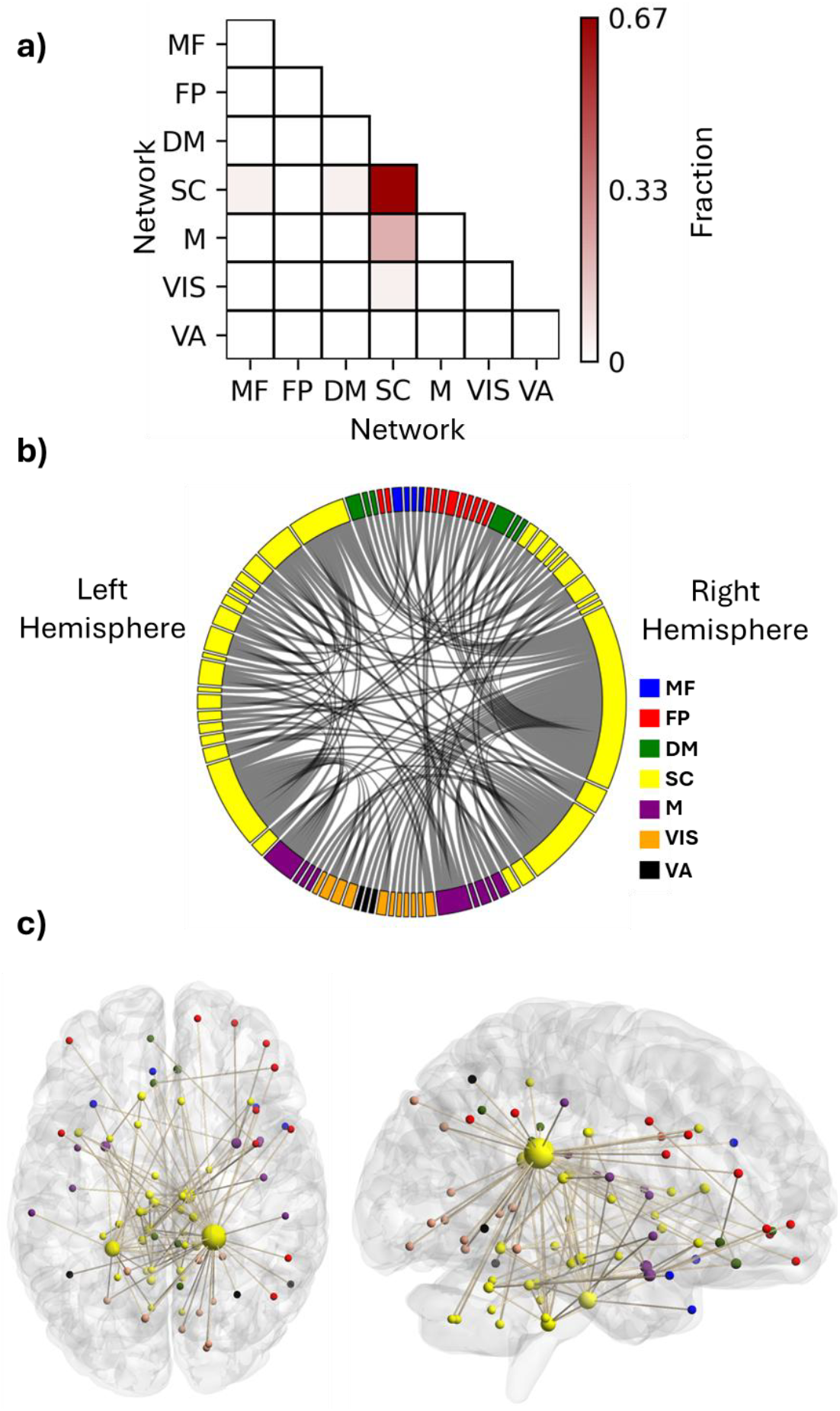
**(a)** Fraction of features with suprathreshold differential power within and between the functional networks. Each element shows what fraction of the discriminative edges resides between the networks. **(b)** distribution of the discriminativeFCs among the seven investigated networks. **(c)** An anatomical visualization of the location of the ROIs containing suprathreshold DP features in the brain. Larger regions show a higher number of differentiating FCs. The list of networks is as follows: MF, the medial frontal network; FP, the frontoparietal network; DM, the default mode network; SC, the subcortical-cerebellum network; M, the motor network; VIS, the visual I and II network; VA, the visual association network.

We selected the top 3% of ROIs based on their suprathreshold DP edge count to identify the nodes that contributed most to identification, all of which were categorized within the SC network. (Table 1). Interestingly, the top two nodes were contralateral ROIs, left and right caudate tail, which had 38 and 18 high-DP connections, respectively (the two largest yellow spheres in Figure 4c). Most of the right caudate tail’s suprathreshold FCs were with ROIs of the visual system networks (VIS and VA) bilaterally and the right frontoparietal network (Figure 4b). The two caudate regions were followed by ROIs from the brainstem and the cerebellum. Specifically, one ROI encompassed the pons in the right hemisphere of the brainstem; another included both the pons and medulla in the left brainstem, and a third spanned the culmen of the left cerebellar vermis (Table 1), all of which contained both inter- and intra-hemispheric edges.

**Table 1.** Highest contributing ROIs to identification. The ROIs of the SC network with the highest numbers of connections with suprathreshold (99.5 percentile) FCs. The MNI coordinates are the means of the locations of all voxels within each ROI.

| Anatomical ROI | # edges | MNI coordinates |
| --- | --- | --- |
| Right caudate tail | 38 | [21.3, -36.4, 22.4] |
| Left caudate tail | 18 | [-23.7, -41.3, 19.9] |
| Right brainstem pons | 17 | [9.9, -18.8, -30.7] |
| Left brainstem medulla/pons | 12 | [-6.9, -33.1, -39.3] |
| Left vermis culmen | 8 | [-4.8, -21.5, -15.9] |

### MVIC prediction

For different values of FST, the correlation coefficient between the predicted and observed MVIC scores was calculated. For all FSTs, the CPM model was able to predict the MVIC scores with significance (p<0.05, Figure 5a and Table 2). For both tasks, the highest correlation coefficient was seen when the FST=0.0005, where approximately 30 highly correlated features passed the threshold in each fold to build the linear model for MVIC prediction. Using this threshold, for the ADT and VDT tasks, the correlation coefficient between the predicted and observed scores was 0.39 and 0.43, respectively (p-value <0.01). It should be noted that for many of the folds there were no features that were positively correlated with the MVIC scores for any of the FST values, and therefore, the models were dominantly made by negatively correlated features.

**Table 2.**
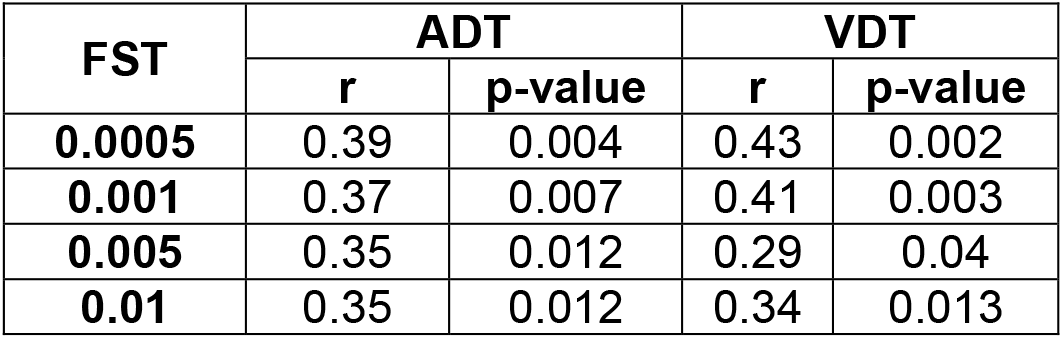
CPM performance in predicting MVIC using ADT and VDT connectomes. The “r” column shows the correlation coefficient between the predicted and observed MVIC scores, and the p-value column shows the corresponding correlation’s statistical significance.

| FST | ADT |  | VDT |  |
| --- | --- | --- | --- | --- |
| | $r$ | $p$-value | $r$ | $p$-value |
| **0.0005** | 0.39 | 0.004 | 0.43 | 0.002 |
| **0.001** | 0.37 | 0.007 | 0.41 | 0.003 |
| **0.005** | 0.35 | 0.012 | 0.29 | 0.04 |
| **0.01** | 0.35 | 0.012 | 0.34 | 0.013 |

**Figure 5.**
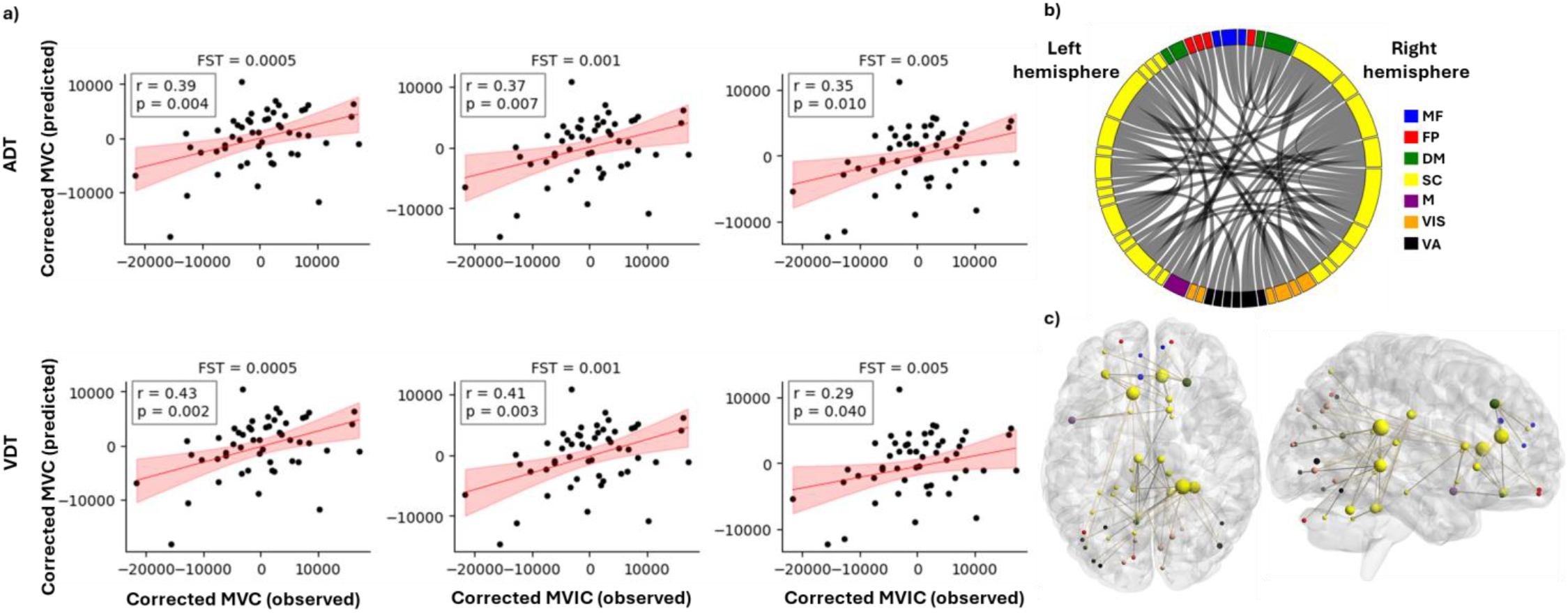
Prediction of MVIC using CPM model. (a) Predicted vs observed MVIC scores after accounting for effects of age and sex for feature selection thresholds of 0.0005, 001 and 0.005 for both ADT and VDT tasks. “r” shows the correlation between the predicted and observed MVIC scores, and “p” shows the corresponding p value of the correlation. (b) distribution of the MVIC predictive FCs among the seven investigated networks. (c) An anatomical visualization of the location of the ROIs with the highest number of MVIC predictive features in the brain. The larger a region is, the higher number of predictive functional connections it has. The list of networks is as follows: MF, the medial frontal network; FP, the frontoparietal network; DM, the default mode network; SC, the subcortical and cerebellum network; M, the motor network; VIS, the visual network; VA, the visual association network.

Depending on the task and FST, a different set of features was used to build the predictive linear model for each fold. To identify features that were robust to outlier participants, we extracted those that were consistently selected across all folds for each task–FST combination (Table 3). In addition, the intersection of the chosen features between the two tasks was examined to ascertain the predictive features that are not specific to either perception task (Table 3). The between-ROI FCs governing the MVIC prediction consistently across the two tasks are shown in Figure 5 b-c, along with their corresponding functional network (FST=0.005). All the features that were common across ADT and VDT were negatively correlated with the MVIC scores.

**Table 3.**
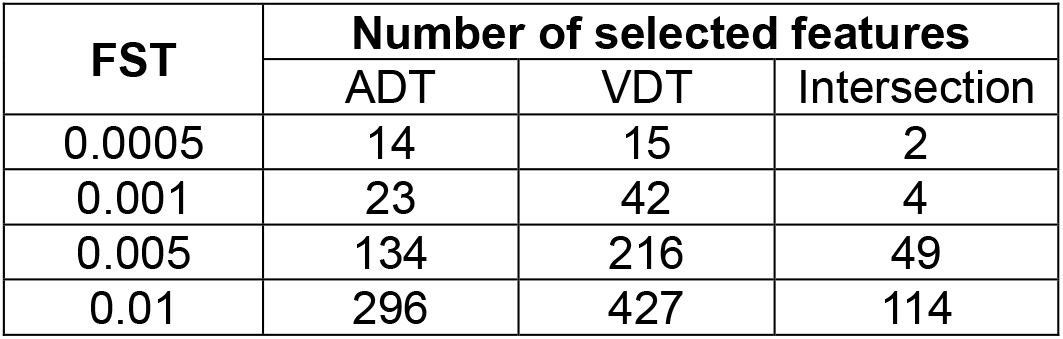
Number of MVIC-predictive functional connections across tasks and FST thresholds. The number of functional connections (FC) correlated with the MVIC scores with a p-value less than the feature selection threshold (FST) are shown for both visual (VDT) and auditory (ADT) tasks. In the intersection column, the number of FCs that had the mentioned property across the tasks are reported.

Following the same approach used for identification, we extracted the top 3% of ROIs with the highest number of MVIC-predictive connections (Table 4). All these nodes were in the SC network, with the right caudate tail— already identified as the most differentiating ROI—contributing the most MVIC-predictive features through 8 functional connections. The other three ROIs that provided the highest number of MVIC predictive FCs were the right cingulate gyrus, the left caudate head, and the right hippocampus tail (Table 4 and the ROIs indicated by the large yellow spheres in Figure 5c). There were also no FCs that appeared both in highly differentiating FCs (suprathreshold DP edges) and the predictive features of MVIC.

**Table 4.**
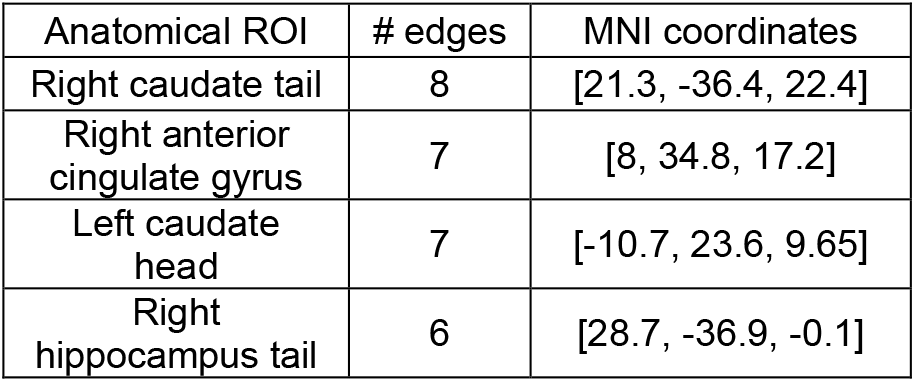
Highest contributing ROIs to MVIC prediction. The ROIs with the highest numbers of connections correlated with MVIC scores.

| Anatomical ROI | # edges | MNI coordinates |
| --- | --- | --- |
| Right caudate tail | 8 | [21.3, -36.4, 22.4] |
| Right anterior cingulate gyrus | 7 | [8, 34.8, 17.2] |
| Left caudate head | 7 | [-10.7, 23.6, 9.65] |
| Right hippocampus tail | 6 | [28.7, -36.9, -0.1] |

In doing the permutation testing, after shuffling the MVIC scores across the participants, for FSTs of 0.0005 and 0.001 we could not perform the permuted predictions, as in some of the folds, there were not any features passing the FST to build the model. Therefore, we performed the permuted predictions using FSTs of 0.005 and 0.01 and found the distribution of correlation between the predicted and observed scores under chance and compared it with the results driven by the unpermuted data. We observed a p-value of 0.03, indicating that the MVIC prediction was unlikely to have occurred by chance.

## 4. Discussion

With increasing life expectancy, frailty has become a widespread public health concern [45], highlighting the urgent need for predictive models at the individual level to enable early detection and targeted interventions. In this study, we employed CPM, a data-driven approach of phenotypic prediction to explain individual differences in MVIC, an index of physical frailty, among healthy older adults using task-based functional connectomes. We showed that connectomes derived from a task with different perceptual but similar motor components captured individual-specific signatures, enabling CPM to predict MVIC. We identified the key networks (intra-SC, SC-M), ROIs (bilateral caudate tail, pons), and FCs (right caudate tail, right anterior cingulate gyrus, right hippocampus tail) that contributed to the identifiability of the connectomes and predictive of MVIC to find the neural substrates of individual-specific levels of frailty. These findings demonstrate CPM’s potential as a predictive model and biomarker for frailty, enabling early identification and personalized intervention strategies.

We observed that the FCs within the SC network and between the SC and M networks had the highest contribution to providing identifiability of the task-based connectomes for older adults. This could stem from the handgrip manipulation common to both tasks. Previously it has been shown that engagement in a motor task would alter the motor network connectivity and increase its inter-individual variability [46]. The motor part of our tasks was a combination of force generation (applying an amount of physical power) and force control (keeping the force in the high grip or low grip range in the squeeze period of the trial) which engages both M and SC networks to coordinate the demands of the task [47]. This being the case, the high engagement of these networks would transfer their connectivity patterns to a task-specific state which makes them highly identifiable. In contrast, studies of resting-state connectomes in younger adults have shown that the majority of discriminative FCs originate from the MF and FP networks [14, 48]. Older adults tend to have high connectivity in their frontal brain regions which could cause a similar connectivity pattern within MF and FP regions across our cohort of older adults [49], therefore, reducing their corresponding FCs differentiation power.

At the ROI level, the bilateral caudate tail areas had the most differentiating FCs across the dataset. The caudate nucleus is one of the subcortical regions known to be correlated with force generation [50], and its functional connectivity pattern would significantly change in the period of force exertion, making its FCs individual-specific. Because the participants squeezed the dynamometer with their non-dominant hand, it may explain why the right caudate nucleus tail was more involved in the force generation and shaped more participant-specific FCs compared to its left counterpart. Following the caudate regions, we identified ROIs in the right brainstem pons and the left brainstem pons extending into the medulla which exhibited the most differentiating FCs. The basal ganglia–brainstem pathway is key to motor control, helping to integrate voluntary and automatic movements. It regulates postural muscle tone and locomotion by sending signals to brainstem motor networks, which are essential for starting and sustaining movement [51]. Given this role, its functional connectivity pattern would be altered during a task with a squeeze component, which could induce a differentiating pattern across the dataset. Furthermore, these brainstem ROIs’ FCs with the right caudate tail were among the features with the highest differential power, showing the contribution of basal ganglia – brainstem connections in individuals’ identifiability. Finally, the culmen area of the vermis of the left cerebellum also had distinct and identifiable FCs. While this area is relatively understudied, its functional connectivity is correlated with gait impairment in PD patients, which shows its potential significance in motor-related activities [52].

In the prediction of MVIC, the ROI corresponding to the right caudate tail had the highest number of FCs correlated with MVIC scores. Previous studies showed that brain perfusion in the caudate nucleus is a strong predictor of frailty, distinguishing between frail, prefrail, and nonfrail HIV patients [53], and its volume reduction is associated with frailty in cognitively impaired older adults [54]. The caudate nucleus and putamen form the neostriatal nucleus, which receives inputs from the cerebral cortex and transmits them to the brainstem and spinal cord, playing a crucial role in the regulation of motor functions [55]. More importantly, in terms of functional connectivity, it was found that that the caudate nucleus, particularly the left side, had the highest number of FCs that were predictive of self-reported frailty index among all brain regions [27]. The right caudate tail and the left caudate head were predictive of MVIC in this study as well, which further supports the potential of this region’s functional connectivity to explain individual differences in frailty. The next ROIs whose FCs were predictive of MVIC scores were the right anterior cingulate gyrus and the right hippocampus tail. While anterior cingulate gyrus is not directly related to force generation, it plays a vital role in decision making and many other cognitive processes [56], which could be involved in performing a task following the prompts given to the participant. Finally, the hippocampus, with its well-established role in memory and processing sensory information [57], seems to be involved in defining physical force and potentially frailty for older adults.

There are some limitations in this study. MVIC is related to various other factors beyond the brain connectome, sex and age variables considered here. These include but are not limited to corresponding muscles’ cross-sectional area [5, 6], fatigue [58], and sleep deprivation [59]. Future work that incorporates these factors may result in a more reliable frailty predictive model. In addition, several cerebellar ROIs were excluded from the analysis because they were not captured within the FoV for this study. These regions, along with their corresponding FCs with other ROIs, may contain predictive features for frailty, potentially enhancing CPM performance in predicting MVIC. Our dataset was moderately sized, which might limit the generalizability and replicability of the MVIC prediction using task-based connectomes. Future research should explore the effects of several factors on MVIC prediction using functional connectome, such as parcellation scheme, preprocessing and standardization framework [60], task paradigm, and the dynamic nature of the brain connectome (i.e., time-varying brain connectivity states) that has been shown to outperform the static connectome in predicting certain behavioral measures [61].

Despite these limitations, this is the first study to show that a motor task connectome is individual-specific and can predict frailty (handgrip strength) in older adults. The SC network contained most of the differentiating and predictive functional connections, highlighting its role in grip strength variability. Notably, the caudate emerged as the most significant region in MVIC prediction, emphasizing the need for further research in its role in frailty.

## 5. Conclusion

In this study, we applied connectome-based predictive modeling on task-based functional connectivity data to predict an index of frailty, handgrip maximum voluntary contraction strength, in older adults. The model demonstrated statistically significant predictive power for individual differences in contraction strength., suggesting that the task-based connectomes can be informative of frailty state for this cohort. The subcortical areas of the brain, specifically the caudate nucleus, played a key role in providing this predictive power for the connectomes. These findings highlight the potential value of subcortical brain regions, particularly the caudate nucleus, as targets for further investigation into the neural basis of frailty and as potential sources for developing targeted interventions or treatments.

## Acknowledgments

This work was supported by R01-AG072607 from the National Institutes of Health/National Institute on Aging.

